# A modified GC-specific MAKER gene annotation method reveals improved and novel gene predictions of high and low GC content in *Oryza sativa*

**DOI:** 10.1101/115345

**Authors:** Megan J. Bowman, Jane A. Pulman, Tiffany L. Liu, Kevin L. Childs

## Abstract

Accurate structural annotation depends on well-trained gene prediction programs. Training data for gene prediction programs are often chosen randomly from a subset of high-quality genes that ideally represent the variation found within a genome. One aspect of gene variation is GC content, which differs across species and is bimodal in grass genomes. We find that gene prediction programs trained on genes with random GC content do not completely predict all grass genes with extreme GC content. We present a new GC-specific MAKER annotation protocol to predict new and improved gene models and assess the biological significance of this method in *Oryza sativa*.

## Background

Most widely used gene prediction programs depend on Hidden Markov Models (HMMs) to predict gene structure within genomic sequence [1–3]. Typically, genes are modeled within HMMs using a series of hidden states that represent generic gene structure. The hidden states are filled with transition probabilities based on k-mer sequences taken from the genes that are used to train the HMM. It is known that gene GC content can affect gene predictions. Korf found that accuracy of predicting genes in one species using a SNAP HMM that was trained for a second species was more correlated with the GC content of the two species’ genomes than with the phylogenetic distance between the two species [1]. Additionally, in mammalian genomes, which contain so-called isochores, gene GC content is correlated with the GC content of the surrounding genome. The AUGUSTUS gene prediction program has a feature that trains multiple HMMs that are each specialized for different narrow isochore-specific GC ranges in order to improve gene predictions [4–6].

We perceived that two factors might limit the accuracy of gene prediction in grass genomes. First, in many species including most plants, the GC content of genes has a relatively narrow and unimodal distribution, but in the grasses (Poaceae), the GC content of genes has a broad bimodal distribution (Figure 1A; [4,7–10]). The bimodal distribution of GC-content in the grasses suggests that there exist two classes of genes (high GC and low GC) that the gene prediction programs are attempting to learn. While gene prediction programs perform well with grasses [11], we hypothesized that the accuracy of grass gene predictions could be improved by accounting for the high and low GC gene classes. Furthermore, as with any supervised machine learning technique, we expect that it is difficult to predict grass genes at the tails of the natural GC distribution and that some grass genes may not be predicted at all using existing protocols. Second, grass gene GC content is not well correlated with the surrounding genomic regions (Figure 1B; [10,12,13]), and therefore, grass genomes do not contain isochores. We also predict that grass genome annotation will not benefit from analysis by the isochore-sensitive AUGUSTUS protocol [6]. Therefore, it is probable that gene annotation in grasses can be improved further.

**Figure 1.**
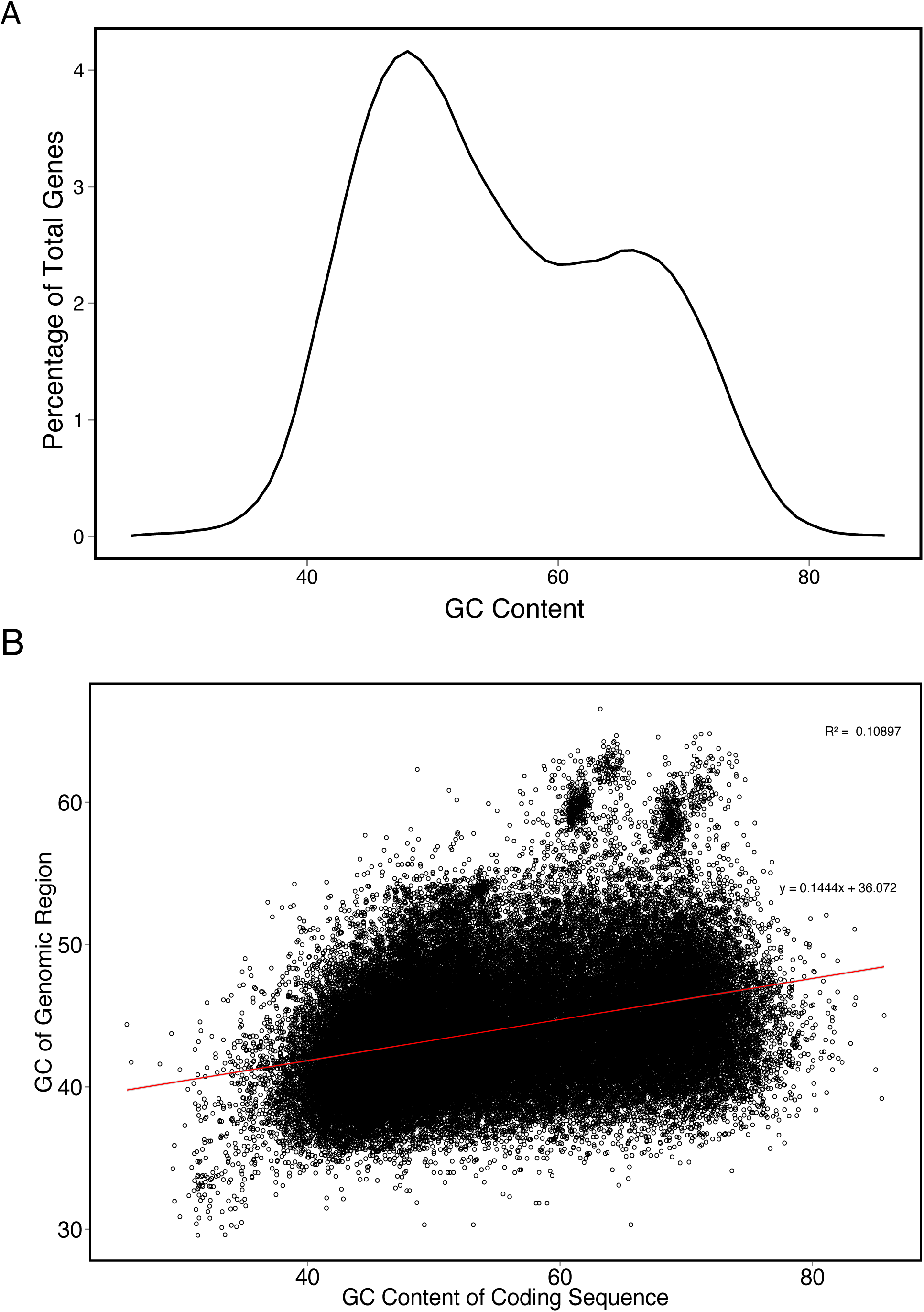
Bimodal distribution and coding region GC content in *Oryza sativa*. A) Distribution of GC content of IRGSP v7 predicted coding regions. B) GC content of IRGSP v7 predicted coding regions vs. genomic GC content 5Kb upstream and downstream of predicted coding regions.

MAKER is a commonly used structural annotation engine that has been used to annotate numerous plant genome assemblies [11,14–18]. The MAKER gene annotation pipeline makes it very easy to train and then predict gene models from commonly used *ab initio* gene prediction programs, such as SNAP and AUGUSTUS [1,6,19]. We developed a new GC-specific MAKER protocol that makes use of genes with high and low GC content as training data in order to derive separate versions of the SNAP and AUGUSTUS HMMs that are tuned to accurately predict high and low GC genes. Using this new method, we improved regular MAKER gene predictions in *Oryza sativa* (rice) relative to available transcript and protein evidence. Furthermore, we identified novel genes with high and low GC content that had not been predicted by the standard MAKER protocol. Comparisons to the AUGUSTUS isochore-based prediction method as well as to the standard MAKER protocol showed that this GC-specific MAKER protocol shifts the overall GC content of predicted gene models both higher and lower than the standard MAKER protocol. This new GC-specific MAKER annotation method will be of interest to anyone working on structural annotation of genomes with bimodal GC content but will likely improve the annotation of any genome.

## Results

### Reannotation of the *O. sativa* genome with MAKER using HMMs trained on high and low GC content

We thought that grass genes identified by gene prediction programs that are trained on genes with specific GC content could both find different genes and produce differing gene models at identical loci than prediction programs that are trained on genes with random GC content. We tested this hypothesis by reannotating the genome of *O. sativa.* In order to compare gene models within the *O. sativa* ssp. Nipponbare genome (v7 assembly; [20]) based on the GC content of different HMM training sets, three MAKER structural annotations were completed using a modified method. SNAP and AUGUSTUS HMMS were trained either with training genes randomly picked without regard for GC content, with training genes with low GC content or with training genes with high GC content. The standard MAKER annotation using HMMs trained on randomly selected training genes for SNAP and AUGUSTUS predicted 29,133 gene models with transcript evidence and/or Pfam protein domains. The structural annotation based on high GC HMMs produced 26,063 evidence supported gene models, and the MAKER annotation based on low GC HMMs produced 26,559 evidence supported models (Table 1). The average length of transcripts was very similar for the standard and low GC structural annotations (Table 1). The average transcript length of the high GC predictions was considerably shorter, a trend that has been previously discussed in eukaryotic genomes [21].

**Table 1.** Numbers of high quality rice genes predicted by different MAKER protocols.

| Annotation | Number of Predictions | Average Transcript Length | AED<sub>0.5</sub> | Percentage (%) |
| --- | --- | --- | --- | --- |
| Standard Protocol | 29133 | 1920 | 26809 | 92.0 |
| High GC | 26063 | 1439 | 22600 | 86.7 |
| Low GC | 26559 | 2046 | 25091 | 94.5 |
| Six HMMs | 29942 | 1947 | 27395 | 91.5 |

The distribution of GC content of the gene predictions varied greatly (Fig. 2). The standard MAKER annotation has a bimodal distribution of gene GC content with a major peak at 49% and a minor peak at 68%. The GC distribution of the high GC annotation has a unimodal distribution with a major peak at 68%. The low GC annotation has a bimodal distribution with peaks at 47% and 67%. Notably, few low GC genes were predicted by the high GC HMMs, and a lower percentage of high GC genes were predicted by the low GC HMMs compared to the standard GC neutral MAKER annotation.

**Figure 2.**
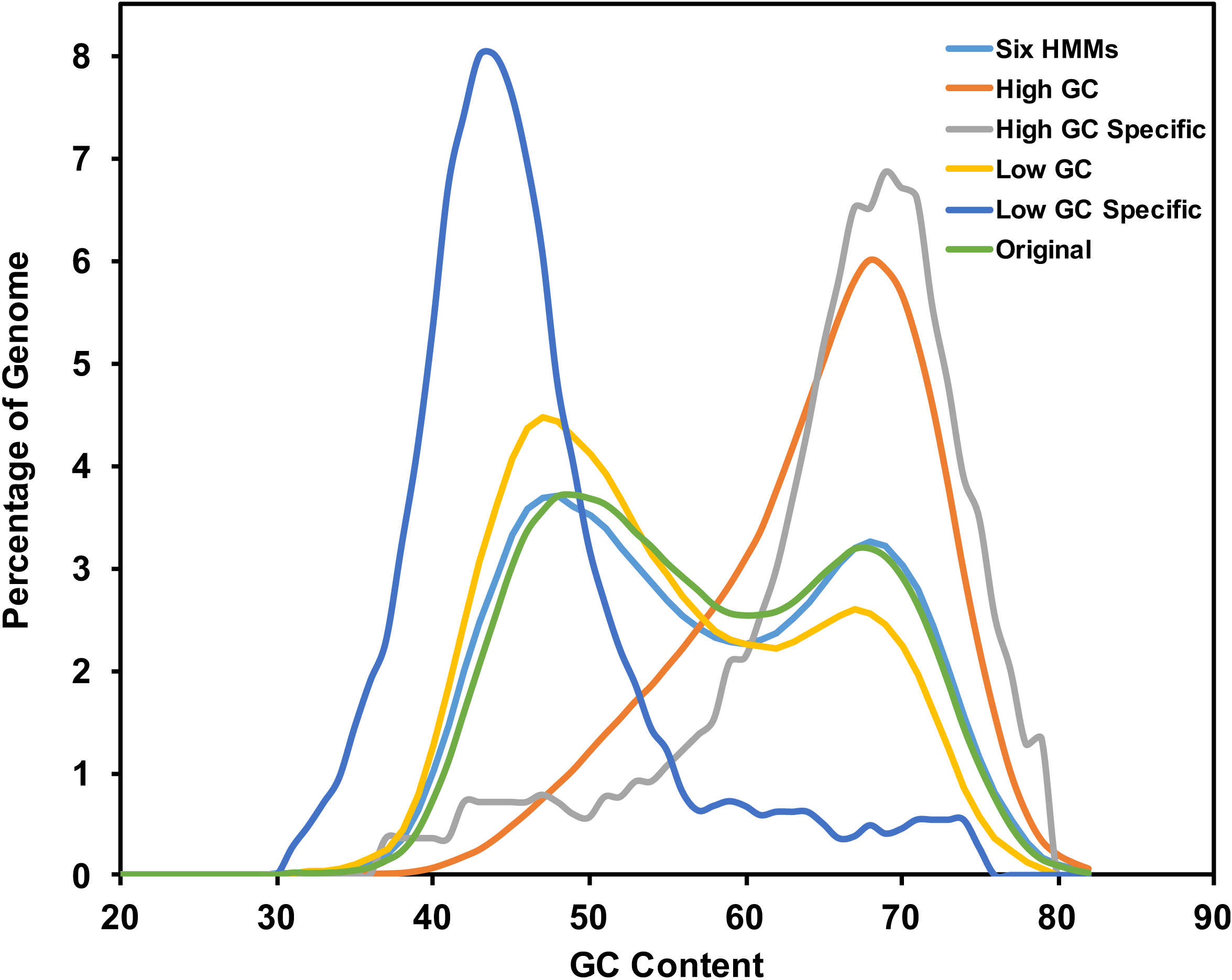
Distribution of GC content of high-quality MAKER gene predictions. Distribution of GC content of various MAKER annotations created through the GC-specific MAKER protocol. The high-quality standard and high GC MAKER genes retain the bimodal distribution that is common to the Poaceae, while the high-quality low GC MAKER genes and the novel high and low GC gene predictions have unimodal distributions centered on GC content associated with the GC content of the HMM training data.

The SNAP and AUGUSTUS HMMs created for the standard, high and low GC MAKER structural annotations were also used together in a single MAKER run to produce a six HMMs annotation (Fig. 3). For this annotation, up to six *ab initio* predictions could be produced at a single locus, but when provided with multiple gene predictions at a single locus, MAKER chooses the single best gene model at that locus. The six HMMs annotation contained 29,942 evidence supported gene predictions (Table 1). The GC distribution for the six HMMs gene set was bimodal with a major peak at 48% and second peak at 68% (Fig. 3). In comparison to the MSU Rice Genome Annotation Project (Release 7) annotation [20], 2,448 gene predictions were unique to the MAKER six HMMs annotation of *O. sativa* while 7,004 gene models found in the MSU annotation were missing from the SixHMMs annotation (Additional File 1).

**Figure 3.**
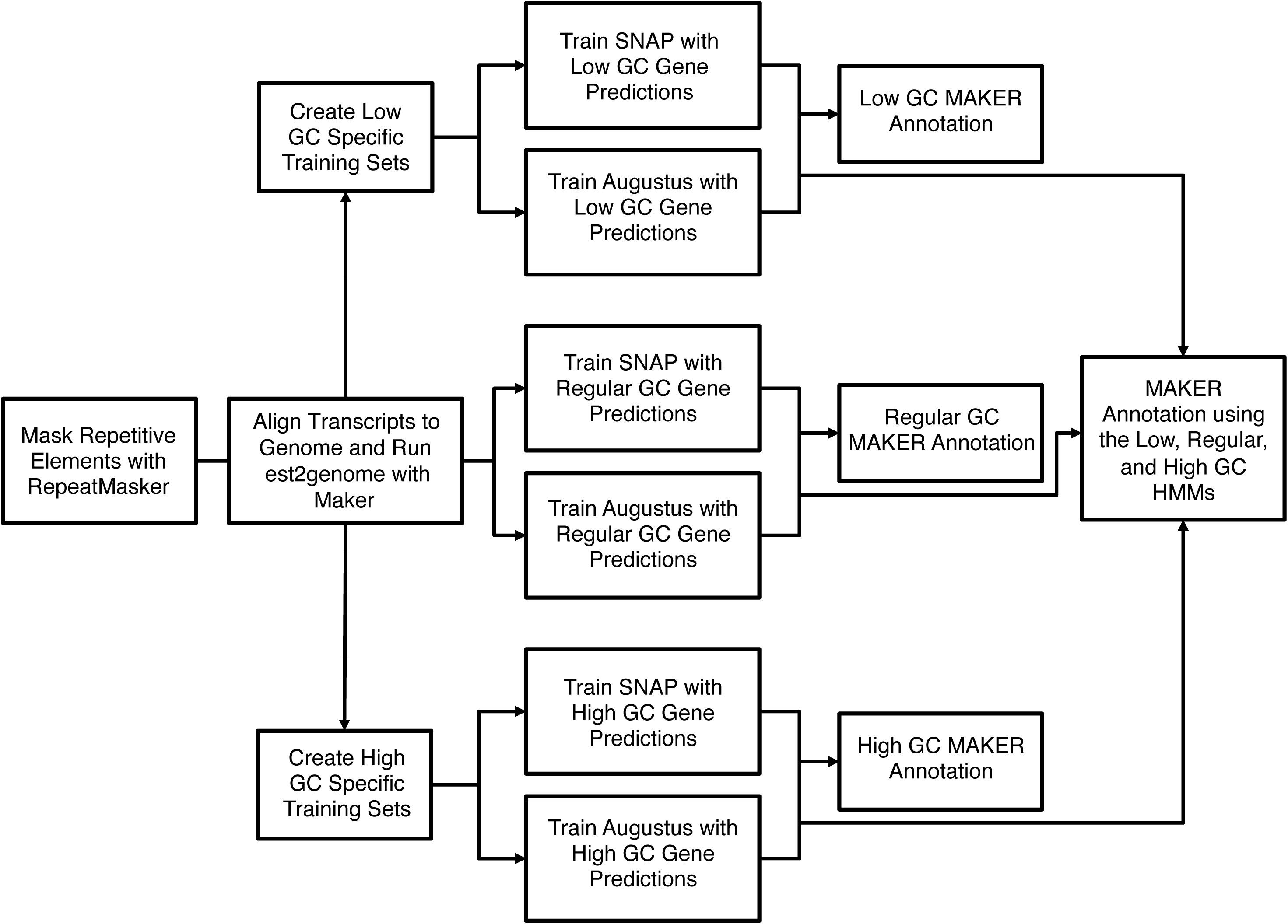
Six HMMs MAKER structural annotation method. The center workflow depicts the standard method for training hidden markov models for use in MAKER, while the low GC (top) and high GC (bottom) training methods can be used after creating high and low GC HMM training data sets. After separately training HMMs with the low and high GC training data, all three SNAP HMMs and all three AUGUSTUS HMMs were specified in the maker_opts.ctl file (see the Methods section), and MAKER was run to create the six HMMs annotation, which incorporates gene predictions from the standard, high and low GC MAKER runs.

To assess the impact of high and low GC specific HMMs on the structural annotation of *O. sativa,* GC content and annotation edit distance (AED) scores were plotted for each set of predicted gene models and visualized as heatmaps (Fig. 4). AED scores are assigned by MAKER and can be used to assess gene prediction quality [22]. AED measures the concordance of a gene prediction relative to the transcript and protein evidence that supports it. AED scores range between 0 and 1, where 0 indicates perfect concordance between the gene prediction and evidence and 1 indicates that no transcript or protein supports the prediction. Genes predicted by HMMs trained on specific GC content caused a general shift in the GC distribution of predicted gene models for both the high and low GC annotations, in comparison to the standard MAKER annotation (Figs. 4A and 4B). In addition to this shift, standard MAKER gene predictions were improved by high or low GC HMMs as determined by a decrease in AED scores between overlapping gene predictions from the standard MAKER and high or low GC HMMs annotations (Fig. 4E, 4F). The number of standard protocol gene models improved in the six HMMs annotation was 3,740. The number or percent of genes with AED scores less than 0.5 (AED_0.5_) can be used for genome wide assessment of annotation quality. The percentages of AED_0.5_ genes were similar for all three annotations (Table 1). The high percentage of well-supported gene predictions reflects the quality of transcriptome evidence provided during the structural annotation process.

**Figure 4.**
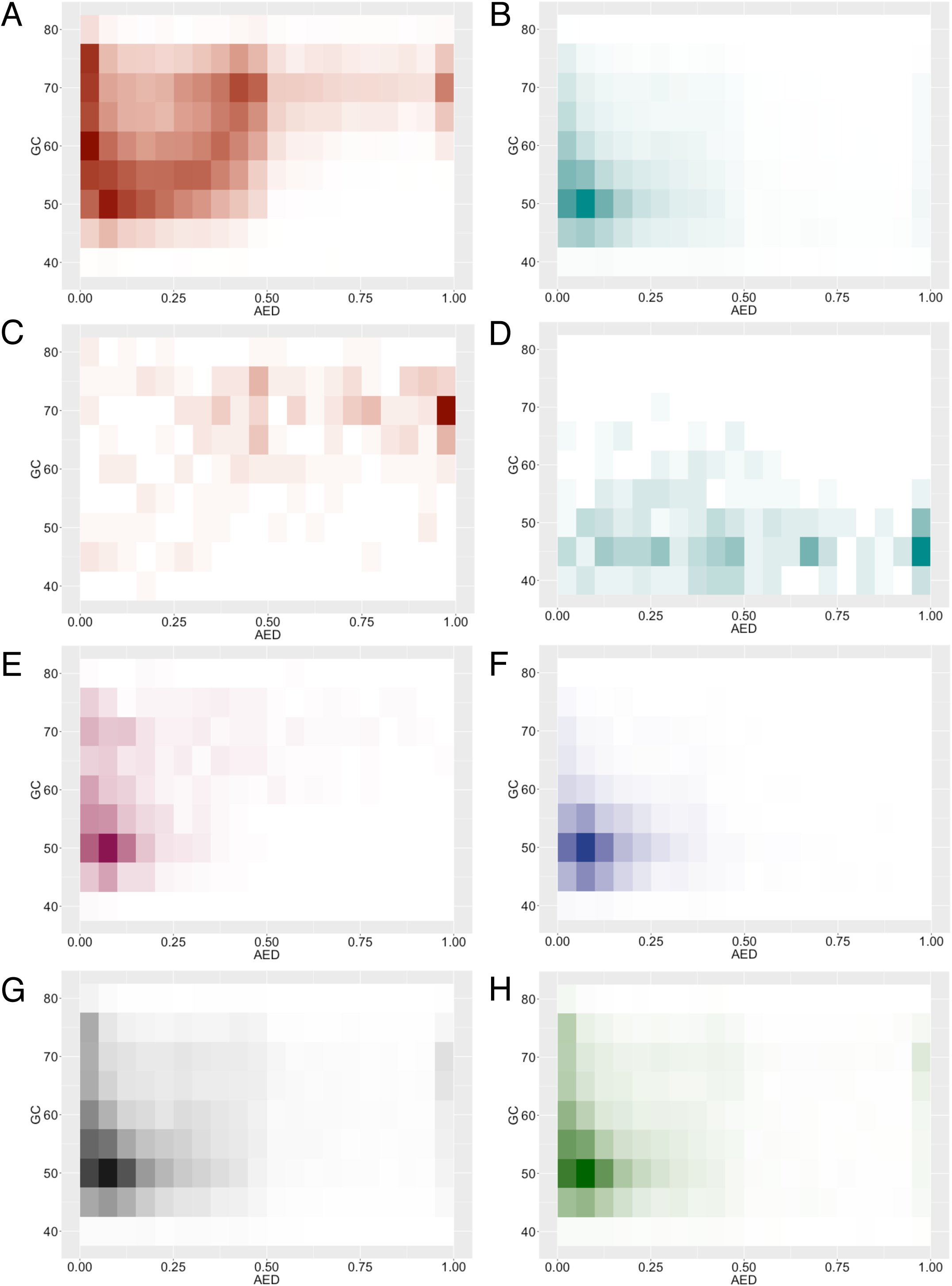
Heatmap visualization of annotated edit distance (AED) and GC content of MAKER predicted gene models. A) MAKER genes predicted using the high GC HMMs. B) MAKER genes predicted using the low GC HMMs. C) Novel genes predicted using the high GC HMMs. D) Novel genes predicted using the low GC HMMs. E) Gene predictions from the high GC HMMs that improved gene predictions made by the standard MAKER protocol. F) Gene predictions from the low GC HMMs that improved gene predictions made by the standard MAKER protocol. G) Gene predictions from the standard MAKER protocol. H) Gene predictions from the MAKER six HMMs annotation.

### Comparison of MAKER six HMMs method to alternative MAKER approaches

The results of the MAKER six HMMs structural annotation were compared to MAKER genome annotations where combinations of the SNAP and AUGUSTUS gene prediction programs were used with alternative parameters. As AUGUSTUS can be run so that it considers GC content of the genomic region (isochores) in which a gene prediction is made, we also trained AUGUSTUS in its isochore-sensitive mode and used it to make gene predictions within MAKER. Overall, the MAKER six HMMs annotation produced more genes than any other annotation strategy tested here (Table 1, Additional File 2: Table S1). MAKER run with only SNAP identified more evidence supported genes than either AUGUSTUS alone trained with randomly chosen training data or the isochore-specific AUGUSTUS protocol. Only a few hundred more genes were generated by the isochore-specific AUGUSTUS annotation than by the randomly trained AUGUSTUS HMM. Using randomly trained SNAP with either randomly trained or isochore-specific AUGUSTUS produced similar numbers of gene predictions but more than when MAKER is run with any of these programs alone. The number of AED_0.5_ gene predictions follows a similar trend to the total number of gene predictions made by any annotation protocol (Table 1; Additional File 2: Table S1; Additional File 3: Fig. S1). However, as more genes are identified by a particular annotation method, the proportion of AED_0.5_ genes decreases. The isochore-specific AUGUSTUS and the randomly trained AUGUSTUS and SNAP gene predictions did not vary in overall GC content (Additional File 3: Fig. S2).

For any machine learning protocol, different sets of training data can lead to slightly different prediction results. To ensure that the results that we observed when we trained SNAP and AUGUSTUS on high and low GC content training data sets were not random, we repeated the standard MAKER annotations three times using independently generated training data. The number of predicted gene models differed by less than 150 in the three randomly replicated standard MAKER annotations (Additional File 2: Table S2), and the AED cumulative frequency plots were nearly identical (Additional File 3: Fig. S3).

### Identification of novel high and low GC content genes

In addition to the improved high and low GC structural annotations created with the MAKER six HMMs annotation protocol, we discovered novel gene predictions specific to the annotations from the high and low GC HMMs. The low GC annotation contained 369 novel genes, while the high GC annotation contained 282 novel genes. Interestingly, the novel genes predicted by the low GC HMMs did not always have a low GC content, and some of the novel genes predicted by the high GC HMMs did not have high GC content (Figs. 2, 4C, 4D). The locations of the novel high and low GC HMM predictions were distributed across all twelve *O. sativa* chromosomes (Table 2; Additional File 4). Of the novel high GC HMM predictions, 253 genes (90%) had some level of protein or transcript evidence for the prediction, while 324 (88%) novel low GC HMM predictions had protein or transcript support (Fig. 5). Overall, the AED scores increased as GC content increased for the novel high GC HMM predictions and as GC content decreased for the novel low GC HMM predictions (Figs. 4C and 4D). The average length of the novel high GC genes was 1,439 bp, while the novel low GC genes were on average 2,046 bp in length. We compared these novel gene predictions to the MSU Rice Genome Annotation Project (MSU-RGAP) Release 7 gene set and found 112 of the low GC HMM predictions and 167 of the high GC HMM predictions were present in that high-quality gene set [20].

**Figure 5.**
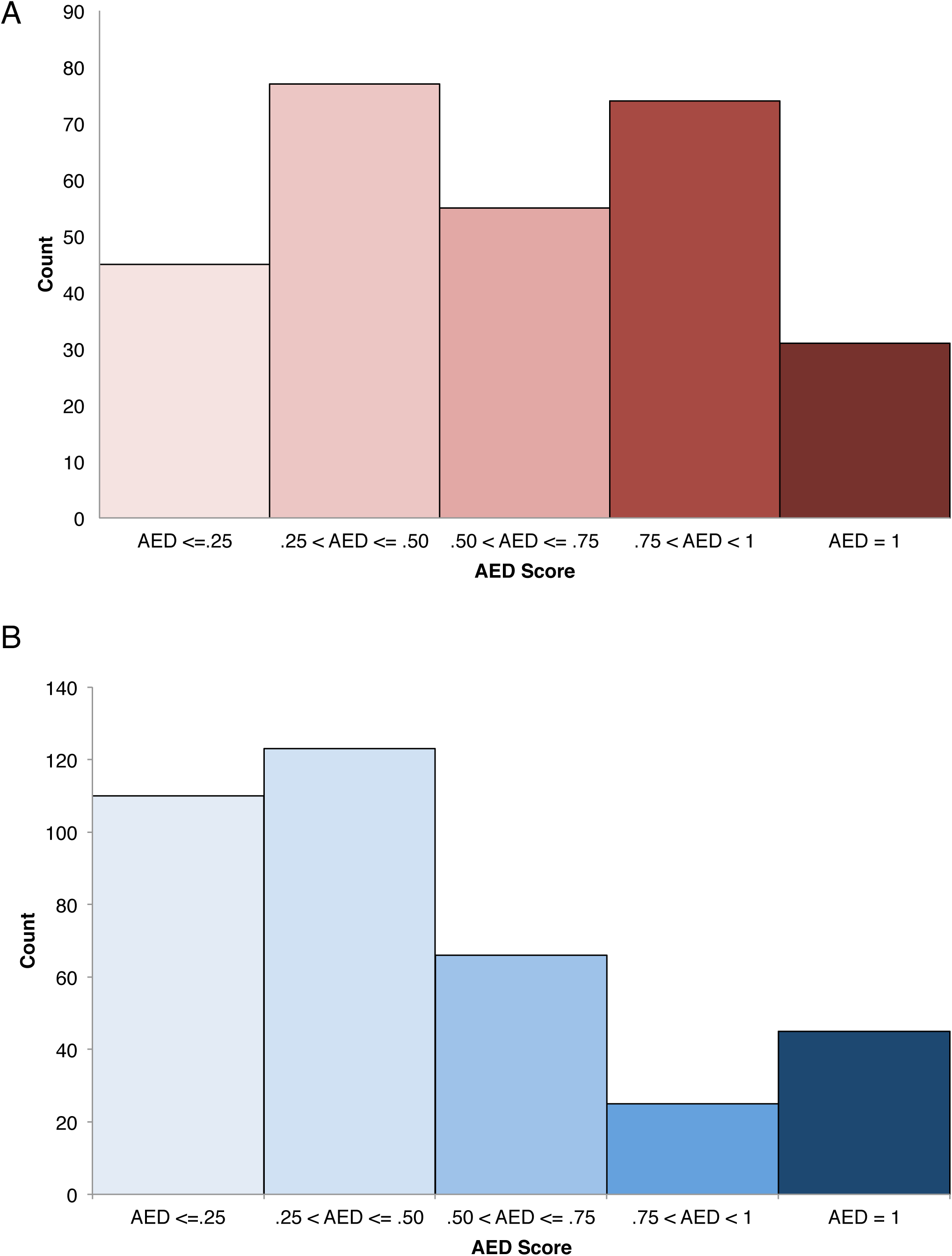
AED scores of high and low GC novel genes in *Oryza sativa*. AED scores for the novel A) high and B) low GC gene predictions generated through the MAKER sixHMM annotation method.

**Table 2.** Distribution across the genome of rice of novel genes predicted by SNAP and AUGUSTUS HMMs trained genes with high and low GC content.

|  | Novel Low GC | Novel High GC<sup>892</sup> |
| --- | --- | --- |
|  | HMM Predictions | HMM Predictions |
| Chr1 | 28 | 33 |
| Chr2 | 39 | 23 |
| Chr3 | 32 | 26 |
| Chr4 | 42 | 35 |
| Chr5 | 35 | 20 |
| Chr6 | 31 | 30 |
| Chr7 | 29 | 18 |
| Chr8 | 26 | 22 |
| Chr9 | 20 | 23 |
| Chr10 | 29 | 21 |
| Chr11 | 31 | 20 |
| Chr12 | 31 | 26 |

### Orthology of high and low GC novel genes to genes from other grass species

Using the total predictions generated through the MAKER six HMMs annotation, additional support was given to the novel predictions made by the high GC and low GC HMMs by first assessing sequence homology of the novel gene predictions to the NCBI non-redundant protein database [23]. Of the 651 novel predictions, 387 had a significant BLASTP hit (e-values less than 1e-10) to NCBI’s non-redundant protein database. Second, the homology and orthology of these genes was evaluated relative to other MAKER six HMM predictions and *Brachypodium distachyon, Sorghum bicolor* and *Zea mays* using OrthoMCL [24–27] (Additional File 5). Of the novel high GC predictions, 51 genes were placed into orthogroups, with 19 as putative homologs only with other MAKER six HMMs predictions, and 32 were orthologous to genes from at least one of the other grass species. Interestingly, 23 novel high GC genes represented the only rice prediction in their orthogroup, and 11 novel high GC genes were single copy orthologs with the other grasses. Of the novel low GC predictions, 92 genes were placed into orthogroups, with 34 as putative homologs only to other MAKER six HMMs gene predictions, and 58 orthologous to the other grass species. Twelve novel low GC predictions were the only rice representatives in their orthogroups.

### Translating Ribosome Affinity Purification (TRAP) sequencing provides additional evidence for novel high and low GC gene predictions

In an effort to demonstrate additional support for the new GC specific gene models outside of the transcript data provided during the MAKER annotation process, translating ribosome affinity purification sequence (TRAP-seq) reads were pseudoaligned to the MAKER six HMMs annotation [28,29], and translatome enrichment indices (TEI) were calculated for each of the novel genes predicted by the high and low GC HMMs. The TRAP-seq samples were isolated from callus, panicle and seedling tissues of an *O. sativa* modified RPL18 transgene [30,31]. TRAP-seq reads were aligned to 200 (71%) of the 282 novel high GC HMM predictions, and 236 (64%) of the 369 novel low GC HMM predictions. This indicated that in addition to the transcript data already aligned to these predictions during annotation, a majority of these novel predictions are in fact being actively transcribed in various tissues from *O. sativa.* The TEI is the ratio of the transcripts per million (TPM) of TRAP-seq to the TPM of mRNA-seq for a specific locus. High TEIs may indicate preferential translation of a transcript, while very low TEIs can be indicative of limited translation [30]. The calculated TEI of each of the novel genes predicted by the high and low GC HMMs that had TRAP-seq pseudoalignments indicates tissue specificity (Fig. 6).

**Figure 6.**
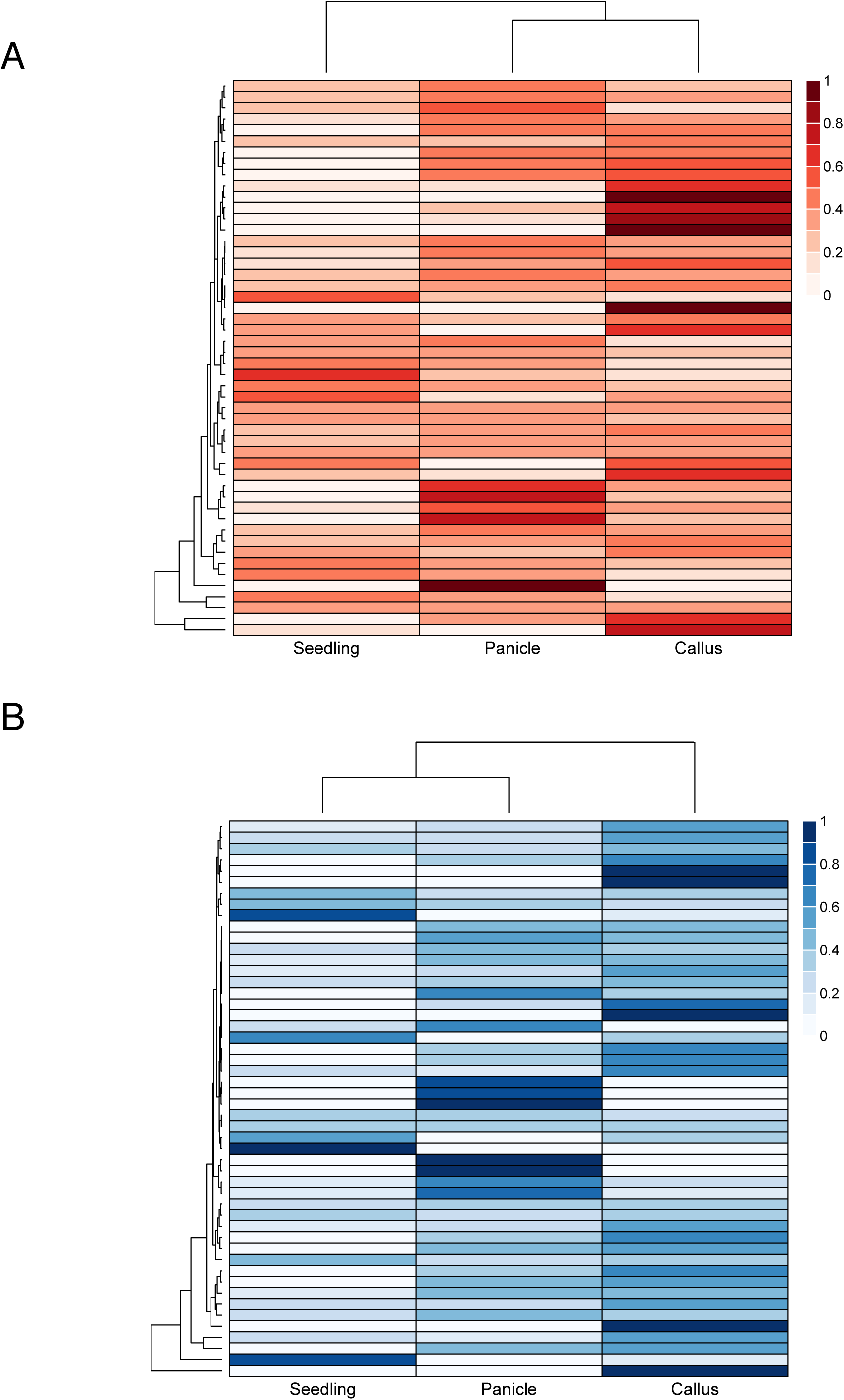
Translatome Enrichment Index (TEI) analysis of novel high and low GC genes. Heatmap of Translatome Enrichment Index (TEI) of the A) novel genes predicted by low GC HMMs and B) novel genes predicted by high GC HMMs gene predictions, which measures the ratio of TRAP-seq to mRNA seq for a specific transcript. Values are scaled by row to a sum of one for visualization purposes.

## Discussion

*Ab initio* gene prediction programs employ HMMs trained on gene sets that should be representative of the variation in gene nucleotide content. We hypothesized that in grass genomes, where genes have a wide variation in GC content and where that distribution is bimodal (Fig. 1A), gene prediction programs trained on random sets of training data would be overly generalized and that this could result in poorly predicted gene models with high or low GC contents. To address this, we developed a GC-specific MAKER gene annotation protocol that trains gene prediction programs SNAP and AUGUSTUS using training data with both high and low GC content. The resulting high-GC and low-GC SNAP and AUGUSTUS HMMs were used in addition to the regularly trained SNAP and AUGUSTUS HMMs to predict genes within MAKER (Fig. 3).

We tested the six HMMs protocol by reannotating the *O. sativa* genome, and we identified 29,942 genes with transcript, protein or Pfam protein domain support. As expected, when MAKER predicted genes in the *O. sativa* genome using either the high-GC or low-GC SNAP and AUGUSTUS HMMs, the GC content of the resulting gene predictions were shifted higher or lower, respectively, compared to the GC content of genes predicted by the standard MAKER protocol (Fig. 2). Furthermore, the GC content distribution of genes predicted by the MAKER six HMMs protocol also showed a shift of the bimodal peaks to higher and lower GC values (Fig. 2). Importantly, most gene predictions made by the MAKER six HMMs annotation overlapped with loci predicted by the standard MAKER protocol, but in 3,740 of these cases, the predictions made by the MAKER six HMMs protocol were improved over the standard MAKER predictions as shown by the better evidence support (i.e. lower AED scores) (Fig. 4E, 4F). This indicates that the high and low GC HMMs were often able to improve upon gene predictions made by the more generally trained gene prediction programs.

In addition to improving the annotation of many genes, we also identified novel genes using this protocol. We found 651 genes that had been identified by high-GC or low-GC SNAP or AUGUSTUS HMMs but that had not been predicted using the standard MAKER pipeline. Of these newly identified genes, 372 were also not found in the most recent MSU-RGAP Release 7 structural annotation [20]. The 279 novel genes predicted by the high-GC or low-GC HMMs that were previously found in the MSU-RGAP Release 7 were likely predicted by MSU-RGAP due to the use of Fgenesh for gene identification, which may have its own biases related to GC content [20,32], or due to the use of different transcript and protein evidence (Additional File 1). Additionally, the MSU-RGAP annotation was improved by PASA, which improves *de novo* gene predictions with transcript alignment evidence, and therefore, PASA is likely not biased by GC content in the same way that HMM-based gene prediction programs can be affected [33]. Furthermore, 90 of the novel genes identified by the high-GC and low-GC HMMs were found to be orthologous to genes from other grass species or to other MAKER six HMMs gene predictions within *O. sativa* (Additional File 5). Additional support for the novel gene predictions comes from examining a TRAP sequencing data set that indicates that 67% of these new predictions are being actively transcribed in three different tissues from *O. sativa* [30] (Figure 6). Nonetheless, as with all computational gene prediction methods, the novel gene models identified by the GC-specific MAKER protocol should be further vetted through additional laboratory analysis.

There are 7,004 genes in the MSU-RGAP Release 7 data set that were not predicted by the six HMMs annotation. Of these genes, 4,327 are characterized as “expressed” meaning that they have may only have transcript support. An additional 1,365 MSU-RGAP genes missing from the six HMMs annotation are described as “hypothetical”, which indicates that they have no transcript or protein support, but they may contain a conserved protein domain (Additional File 1). The expressed and hypothetical MSU-RGAP genes are the genes with the weakest support from that annotation project. Some of the MSU-RGAP hypothetical genes may not pass the stringent evidence test that was applied to the MAKER six HMMs gene predictions which all had transcript or protein support or contained a Pfam domain. Additionally, the MAKER six HMMs annotation only used transcript evidence derived from StringTie assemblies of a small set of RNA-seq reads, but the MSU-RGAP annotation made use of EST and FL-cDNA sequences that were not used in this report. This difference in evidence will have an effect on the genes predicted by MAKER [11]. Finally, the MAKER six HMMs annotation was filtered to remove any predictions that had homology to known transposable elements (TE) and Pfam domains. The MSU-RGAP genes were also filtered to flag any genes with matches to a library of TE sequences, but these two methods were necessarily different and could have resulted in the removal of different subsets of TE-related gene predictions. All of these reasons can help to explain why 7,004 MSU-RGAP genes are not present in our six HMMs MAKER annotation.

Interestingly, there may be additional unrecognized parameters that could be used to improve gene prediction besides our strategy of training gene prediction HMMs in a GC-specific fashion. In the six HMMs annotation, some low GC predictions were generated by the high GC HMMs, and some high GC predictions came from the low GC HMMs (Fig 4A, 4B). While these could be cases of identical gene models being created by two or more HMMs at a particular locus with MAKER randomly retaining only one prediction as the final model for the locus, we also observed novel low GC predictions created by high GC HMMs as well as novel high GC predictions arising from low GC HMMs (Figs. 3, 4C, 4D). This suggests that some unrecognized gene features besides simple GC content were present in the high and low GC HMMs that allowed the prediction of novel low and high GC genes, respectively.

It has been known that the GC content of genes used to train gene prediction HMMs can affect the accuracy of gene predictions [1,6]. The AUGUSTUS gene finder has an isochore-sensitive protocol that was developed in order to more accurately predict mammalian genes. Despite the fact that isochores do not exist in plants (Fig. 1; [12,13]), we used the isochore-sensitive AUGUSTUS protocol to predict genes in *O. sativa,* but we did not see a substantial difference in the number or quality of predicted gene models or a change in overall GC content distribution of those gene predictions (Additional File 3: Figs. S1, S2). This result was expected as gene GC content is not well correlated with the GC content of the surrounding genomic region, and therefore, partitioning the training data before training the gene prediction programs was found to be more effective at improving gene annotations in *O. sativa.*

Given the importance of accurate gene prediction to downstream genomics applications, the GC-specific MAKER protocol described here will be of use to those working on the structural annotation of any species with a bimodal distribution of GC content. MAKER is a powerful tool that enables research groups of any size to pursue structural annotation of sequenced genomes and, with the addition of this protocol, will aid in more accurate gene prediction.

## Conclusions

In this paper we presented a new GC-specific MAKER annotation protocol that was used to successfully identify new evidence supported gene models in *Oryza sativa* with high and low GC content. This new method also improved 13% of gene models produced by the standard MAKER protocol. Comparisons of this method to the standard training protocols for the SNAP and AUGUSTUS *ab initio* gene prediction programs as well as the isochore-sensitive AUGUSTUS gene prediction method showed that by training gene prediction HMMs with data representing multiple ranges of GC content and allowing MAKER to pick the best *ab initio* gene prediction generated by multiple gene prediction HMMs, it is possible to create a final gene annotation set that includes large numbers of both improved and novel gene predictions. The novel gene predictions are supported by various forms of evidence including transcript and protein alignments and membership in ortholog groups with genes from other grass species. Additionally, TRAP-sequencing has shown that a majority of these new predictions are being actively transcribed in *O. sativa.* MAKER is a widely used structural annotation program that allows researchers to produce quality genome annotations. This new method will be an important addition to those interested in the prediction of genes in regions of extreme GC content in Poaceae genomes but will probably be generally applicable for species with narrow, unimodal gene GC distributions as well.

## Methods

### Processing, quality assessment and assembly of evidence

Thirty-one paired end RNA-seq datasets for *O. sativa* grown from different stress environments and tissues were downloaded from the National Center for Biotechnology Information Sequence Read Archive (NCBI-SRA) (Additional File 2: Table S3) using SRAToolkit v. 2.3.4-2 [34]. Raw read quality was assessed with FastQC v. 0.10.1 and Illumina adapters were trimmed using Trimmomatic v. 0.32. Transcripts were assembled using StringTie v. 1.3.0, and these transcript assemblies were subsequently used as EST evidence for all MAKER runs. The SwissProt plant protein dataset was downloaded (ftp://ftp.uniprot.org/pub/databases/uniprot/current_release/knowledgebase/taxonomic_divisions/uniprot_sprot_plants.dat.gz), and all *O. sativa* protein sequences were removed. The remaining protein sequences not from *O. sativa* were used as protein evidence during the MAKER annotation.

### MAKER standard *de novo* structural annotation of *O. sativa*

The MAKER-P (r1128) genome annotation pipeline was used to annotate the Os Nipponbare-Reference-IGRSP-1.0 v7 genome assembly. A custom repeat library was created for *O. sativa* using a method described previously [18]; (http://weatherby.genetics.utah.edu/MAKER/wiki/index.php/Repeat_Library_Construction-Advanced), and the custom repeat library was used by RepeatMasker within the MAKER pipeline to mask repetitive elements. Transcript assemblies and protein sequences described above were used as evidence to aid gene predictions.

A complete description for running MAKER has been provided previously [19,35] and that protocol provides details about ancillary scripts and example command calls. An abbreviated description of the standard MAKER pipeline is given here, and details about the extended GC-specific MAKER pipeline are given below. As MAKER is run iteratively, repeat masking and evidence alignment was performed during an initial MAKER run, and the resulting GFF3 file containing masked regions and protein and transcript alignments was used during all subsequent MAKER runs. The initial MAKER run generates data that aids in the training of the gene predictions programs SNAP (version 2013-11-29) [1] and AUGUSTUS (version 2.6.1) [6] (Figure 3). During the initial MAKER run, the parameter `est2genome` was used to cause MAKER to promote transcript alignments to gene models. High-quality transcript-derived gene models (AED <= 0.2) were used to train SNAP and AUGUSTUS. Instructions for training SNAP can be found elsewhere [1,19,36]. We use a custom shell script, train_augustus.sh, which trains the AUGUSTUS HMM in only a few hours for most species.

train_augustus.sh <path to working directory for training> <path to MAKER gff3 output from initial MAKER run> <species name for AUGUSTUS HMM directory> <path to single fasta file with all transcript assemblies>

The train_augustus.sh shell script prepares training and testing data sets and makes use of the autoAug.pl training script from AUGUSTUS to create the appropriate HMM files. This training script is relatively fast, as it only requires the transcript evidence to be aligned to the genomic regions that contain training and testing gene models instead of aligning those sequences to the entire genome. The working directory is used for writing a number of intermediate files and directories during the AUGUSTUS training process. All transcript sequences that were used as evidence during the initial MAKER run must be placed into a single transcript fasta file and provided here as those sequences will be used during the AUGUSTUS HMM training. The species name provided for the HMM training will be used to name the directory that holds all of the files for the new HMM and is also used to specify the AUGUSTUS HMM in the maker_opts.ctl file. It is necessary to have write permissions in the /config/species directory within AUGUSTUS installation directory in order for this script to work as that is where the AUGUSTUS writes the species-specific HMM directory. On a shared compute system, it may be necessary to make a local installation of AUGUSTUS and to then point MAKER to that installation by updating the path in the maker_exe.ctl file. After training SNAP and AUGUSTUS HMMs, MAKER was then run one last time using only the SNAP and AUGUSTUS HMMs to predict genes. During the final MAKER run, the parameters `keep_preds` was set to 1.

To identify the high-quality gene set, the MAKER accessory scripts gff_merge and fasta_merge, which are included in the MAKER installation, were used to generate a GFF3 file with all gene predictions and evidence data and the transcript and protein fasta files for those predictions. Pfam domains were identified within the predicted proteins using hmmscan[36].

`hmmscan --domE 1e-5 -E 1e-5 --tblout <MAKER max predictions hmmscan output file> <path to Pfam-A.hmm> <path to predicted protein fasta file>`

The annotation GFF3 file, the transcript and protein fasta files and the hmmscan results file were used to generate a quality MAKER standard gene set.

`generate_maker_standard_gene_list.pl --input_gff <output of gff3_merge> --pfam_results <hmmscan output> --pfam_cutoff 1e-10 --output_file <path to MAKER standard gene list>`

`get_subset_of_fastas.pl -l <path to MAKER standard gene list> -f <fasta_merge output transcript/protein fasta> -o <path MAKER standard transcript/protein fasta>`

`create_maker_standard_gff.pl --input_gff <output of gff3_merge> --output_gff <path to MAKER standard GFF3> --maker_standard_gene_list <path to MAKER standard gene list>`

Despite our use of a custom repeat library that was used for masking repeat elements in the genome, some TE-related genes remain unmasked, and we performed additional analyses to identify and remove any TE-related predictions from our MAKER standard gene set. Predicted proteins were compared to a database of Gypsy transposable elements (3.1.b2) [37]. Predicted proteins were also aligned with blastp to a database of transposases [38,39] (http://weatherby.genetics.utah.edu/MAKER/wiki/index.php/RepeatLibraryConstruction-Advanced). A GFF3 file of TE-related genes was derived from the MSU-RGAP gene annotation GFF3 file (http://rice.plantbiology.msu.edu/pub/data/Eukaryotic_Projects/o_sativa/annotation_dbs/pseudomolecules/version_7.0/all.dir/all.gff3) and was compared to the MAKER standard GFF3 file using gffcompare [40].

`hmmscan --tblout <Gypsy HMM analysis output> -E 1e-5 --domE 1e-5 <path to gypsy_db_3.1b2.hmm> <path to maker standard proteins fasta>`

`blastp -db <Tpases020812 database> -query <path to MAKER standard protein fasta> -out <path to Tpases blast output> -evalue 1e-10 -outfmt 6`

`gffcompare -o <TE comparison output file> -r <MSU RGAP TE GFF3> <MAKER standard GFF3>`

The create_no_TE_genelist.py script use the data derived above, the Pfam hmmscan results file and a list of TE-related Pfam domains (TE_Pfam_domains.txt; available on Childs Lab GitHub repository) to create a list of MAKER standard genes with no TE-related predictions.

create_no_TE_genelist.py --input_file_TEpfam <TE_Pfam_domains.txt> --input_file_maxPfam <MAKER max predictions hmmscan output file> --input_file_geneList_toKeep <path to MAKER standard gene list> --input_file_TEhmm <Gypsy HMM analysis output> --input_file_TEblast <path to Tpases blast output> --input_file_TErefmap <TE comparison output refmap file> --output_file <path to TE filtered MAKER standard gene list>

`create_maker_standard_gff.pl --input_gff <MAKER standard GFF3> --output_gff <TE filtered MAKER standard GFF3> --maker_standard_gene_list <path to TE filtered MAKER standard gene list>`

`get_subset_of_fastas.pl -l <path to TE filtered MAKER standard gene list> -f <fasta_merge output transcript/protein fasta> -o <TE filtered MAKER standard transcript/protein fasta>`

This high-quality gene set without TE-related genes was used for all analyses presented in the Results section. In addition to this standard MAKER annotation, two additional annotations were created using either the SNAP HMM alone or the AUGUSTUS HMM alone, and high-quality gene sets without TE-related genes were identified for each of these annotations, which were used for comparisons to the final GC-specific six HMMs annotation described below.

### Training GC-specific HMMs with high-GC and low-GC gene sequences

In order to train high and low GC-specific HMMs for SNAP and AUGUSTUS, it was necessary to use training data that consisted of gene models with CDS GC content within specific ranges. The transcript-based gene predictions from the initial MAKER run (when the `est2genome` parameter was used) served as the starting point for GC-specific HMM training (Fig. 3). After generating the GFF3 file describing the transcript-based gene models, the genome FASTA file was processed by the Perl script MAKER_GC_cutoff_determinati on.pl.

MAKER_GC_cutoff_determination.pl --fasta <full path to file with genome FASTA sequences> --gff full path to MAKER created GFF3 of est2genome> --name <BASE_NAME for output files> --peak <peak determination window, odd integer, default is 5> --smooth <smoothing window, odd integer, default is 7>

The MAKER_GC_cutoff_determination.pl script helps to identify the GC values of the peaks in a bimodal grass gene GC content distribution. The script pulls out the CDS FASTA sequences for the transcript-based gene predictions from the GFF3 and calculates the GC content for each gene prediction. The script assigns the gene GC values to integer bins based on the `--smooth` parameter, which helps to smooth the calculation of the distribution by using a moving window average and writes the results to a file. This output file can be used in R to plot the distribution of gene GC content. A FASTA file of the CDS sequences and a GC content file (showing nucleotide composition and GC content of each prediction) are also produced. In addition, a text file is created with the high and low peak values of the bimodal gene GC distribution that were used here as set points in creating the high and low GC HMM training sets. These peaks are determined by taking each GC bin and looking at a set number of bins on either side (set by `--peak`). A peak is identified when the (`(peak - 1) / 2`) bins on each side of a GC bin have lower calculated GC values than the middle GC bin. However, users may pick their own high-GC and low-GC cutoff values, and the gene GC content distribution graph may aid in picking those cutoff values. The MAKER_GC_training_set_create.py script relies on two files produced from the MAKER_GC_cutoff_determination.pl script: BASE_NAME_gc_content.txt and BASE_NAME_cutoff.txt. The MAKER_GC_training_set_create.py script will create high-GC and low-GC GFF3 files that can be used for training SNAP and AUGUSTUS.

`MAKER_GC_training_set_create.py --input_file_gff <path to MAKER GFF file> --input_file_GC_content <BASE_NAME_gc_content.txt file> --input_file_GC_cutoff`

`<BASE_NAME_cutoff.txt file> --output_file_low <path to the low GC GFF file> --output_file_high <path to the high GC GFF file> --genome_fasta <path to the genome fasta file>`

As detailed in Figure 3, this script is run after est2genome transcript alignment to create new GC-specific HMM training sets. All subsequent steps should only use the high or low GC output files.

### MAKER six HMMs annotation

After the creation of high and low GC SNAP and AUGUSTUS HMMs, a final MAKER run is performed using the standard, high and low GC HMMs at the same time. When using multiple SNAP and AUGUSTUS HMMs for this six HMMs annotation, predictions from the different HMMs can be identified by providing the path to a specific HMM, a colon, and an HMM-specific identifier (see below). Providing a comma-separated list to the `snaphmm` and `augustus_species` parameters allows the designation of multiple HMMs. To create this new six HMMs structural annotation, the following parameters are set in the MAKER maker_opts.ctl file:

`# ----- Re-annotation Using MAKER Derived GFF3`

`maker_gff = path to MAKER alignment GFF3`

`est_pass=1`

`protein_pass=1`

`rm_pass=1`

`# ----- Gene Prediction`

`snaphmm= path to standard SNAP HMM:orig_snap, path to high`

`GC HMM:high_snap, path to low GC HMM:low_snap`

`AUGUSTUS_species= path to standard AUGUSTUS`

`directory:orig_aug, path to high GC AUGUSTUS`

`directory:high_aug, path to low GC AUGUSTUS`

`directory:low_aug`

`keep_preds=1`

Once the six HMMs MAKER annotation is finished, a final high-quality MAKER gene set composed of gene models with transcript, protein or Pfam domain support was created using the same protocol that was used above for the standard annotation.

### Creation of SNAP and AUGUSTUS HMMs trained with transcripts of randomized GC content

To assess the impact of GC specific HMM training on the structural annotation of *O. sativa,* three MAKER annotations were created using HMMs trained with transcripts with randomized GC content from the standard annotation. The following Perl scripts were used, which create the training GFF3 files based on a random seed instead of percentage GC content. The random_dataset_generate.pl script takes as input the MAKER standard transcript FASTA and outputs three transcript FASTAs to be used for downstream GFF3 creation and HMM training.

`random_dataset_generate.pl --transcript <name of transcript FASTA file > --random1 <name of output random file1> --random2 <name of output random file2> --random3 <name of output random file3>`

The seq_name.pl script was run for each of the three random output FASTA files, and generates a list of MAKER standard transcript names from each transcript FASTA.

`seq_name.pl`

`--fastafile <path to an output file from random_dataset_generate.pl>`

`--output <name of text file with ID names for each FASTA sequence>`

Finally, the random`_gff3_create.pl` script requires as inputs the MAKER standard GFF3 with the genome FASTA included and the gene IDs from each of the random FASTAs, and the script generates the final randomized GFF3s that were used for SNAP and AUGUSTUS HMM training.

`random_gff3_create.pl`

`--align_gff <path to MAKER GFF3 with FASTA included>`

`--rand_1 <path to random 1 IDs>`

`--rand_2 <path to random 2 IDs>`

`--rand_3 <path to random 3 IDs>`

The outputs of these steps are three GFF3 files containing the coordinates of randomly selected gene predictions. Each of the GFF3 files created by random_gff3_create.pl was then used for SNAP and AUGUSTUS training.

### Isochore-specific AUGUSTUS training in *O. sativa*

To compare the MAKER GC specific HMM training protocol to the isochore specific AUGUSTUS method, we trained AUGUSTUS in its isochore-sensitive mode as detailed below [6]. After isochore-specific AUGUSTUS training, the resulting HMM was used in a MAKER run with all other parameters as had been used for the standard annotation to create a MAKER structural annotation based only on isochore-specific AUGUSTUS gene predictions. An additional annotation was also created with the isochore-specific AUGUSTUS HMM and the standard SNAP HMM for comparison to the six HMMs annotation method.

After one round of traditional AUGUSTUS training [6] which creates the augustus.gb.train and augustus.gb.test genbank formatted gene files, change the gc_range_min value to 0.32, gc_range_max value to 0.73 and the decomp_num_steps value to 7 in the parameters.cfg file in the newly created AUGUSTUS species HMM directory. The following three commands will then complete the isochore-specific training of AUGUSTUS.

`[AUGUSTUS_installation_dir]/scripts/optimize_augustus.pl --species=<species_name> augustus.gb.train`

`etraining --species=<species_name> augustus.gb.train`

`augustus --species=<species_name> augustus.gb.test`

### Identification of novel high or low GC content gene predictions

The BEDtools v 2.23.0 [41] intersect command was used to compare two GFF3 files containing MAKER gene coordinates to identify novel gene models that were unique to the gene predictions created with high or low GC HMMs. Those predictions that were only created by high GC HMMs but not standard or low GC HMMs were considered novel high GC HMM predictions, while predictions created only by the low GC HMMs were deemed novel low GC HMM predictions.

### Identification of orthologs of novel *O. sativa* gene predictions in other grass species

Paralogs of novel high or low GC gene predictions in *O. sativa* and orthologs in other grass species were identified using OrthoMCL (v1.4) [27] using default parameters with the predicted proteins of the high or low GC unique genes. Predicted proteins of *Brachypodium distachyon* (v 3.1; https://phytozome.jgi.doe.gov/pz/portal.html#!info?alias=Org_Bdistachyon) *Zea mays* (v5b+, Phytozome 11: http://phytozome.jgi.doe.gov/pz/portal.html#!info?alias=Org_Zmays), *Oryza sativa* ssp. Nipponbare (v7.0, http://rice.plantbiology.msu.edu/) and *Sorghum bicolor* (v3.1, https://phytozome.jgi.doe.gov/pz/portal.html#!info?alias=Org_Sbicolor) were used for comparison.

### Translating ribosome affinity purification (TRAP) sequencing analysis

Paired end TRAP-seq and mRNA-seq reads were trimmed using Cutadapt v1.8.1 using default parameters. RNA quantification was conducted with the kallisto v 0.42.5 [42] pseudoalignment method using the six HMMs MAKER predicted transcripts with a bootstrap value of 100. Transcripts per million (TPM) were calculated for both the TRAP-seq and mRNA-seq reads for each tissue, and the translatome enrichment index (TEI) was calculated as the ratio of transcripts per million of TRAP-seq to mRNA-seq for each high-quality six HMMs MAKER transcript. TRAP-seq and mRNA-seq data are available in NCBI BioProject PRJNA298638.

### Figure Creation

Figures were created in R (v. 3.1.1) [43] using the following packages: ggplot2 [44], reshape2 [45] and NMF [46].

## Availability of Data

Perl and python scripts for the GC-specific MAKER protocol are deposited in Github: https://github.com/Childs-Lab/GC_specific_MAKER. Additional relevant data, including FASTAs of MAKER predictions and GFF3 files can be found at the Dryad data repository: (http://www.datadryad.org. Data can only be deposited after acceptance. During the revision process the Dryad DOI will be added here.)

## Acknowledgments

This research was supported by grant IOS–1126998 to KLC from the U.S. National Science Foundation. We would like to thank John Hamilton for help creating the TE-related Pfam domain list.

## Additional Files

**Additional File 1:** (.pdf) **Venn diagram depicting the overlap between the rice GC specific sixHMM annotation and IGRSP v7 annotation**. Of the 7,004 genes that are only present in the IGRSPv7 annotation, 1365 (19.5%) are designated as “hypothetical”, while 4327 (61.8%) are designated as “expressed”.

**Additional File 2**: (.xlsx) **Transcript evidence used to reannotate *Oryza sativa* and additional information from gene predictions generated from alternative MAKER methods**. Table S1. Number of predictions, average transcript length and AED_0.5_ of gene predictions generated by alternative MAKER approaches. Table S2. Number of predictions, average transcript length and AED_0.5_ of the three randomly replicated standard MAKER annotations. Table S3. RNA-seq transcript evidence used in the reannotation of the *Oryza sativa* genome.

**Additional File 3:** (.pdf) **AED curves from various MAKER annotation methods**. Figure S1. AED curves of MAKER annotations of *Oryza sativa* using various *ab initio* prediction methods. Figure S2. Distribution of GC content of MAKER annotations of *Oryza sativa* using various *ab initio* prediction methods. Figure S3. AED curves of MAKER annotations of *Oryza sativa* using HMMs trained with randomized training data.

**Additional File 4:** (.pdf) **Distribution of GC content, MAKER six HMMs gene predictions and novel genes predicted by the high and low GC HMMs in the *Oryza sativa* genome**. A) Genomic GC content in 300Mb bins. Warmer colors indicate higher than average GC content while cooler colors indicate lower than average GC content. B) Heatmap visualization of the density of MAKER six HMMs gene models. C) Genomic location of novel genes predicted by the high GC HMMs. D) Genomic location of novel genes predicted by the low GC HMMs.

**Additional File 5**: (.txt) **OrthoMCL orthogroups containing novel high and low GC gene predictions**. OrthoMCL output listing the number of genes, taxa and gene names for each orthogroup that contains at least one novel high or low GC prediction. Novel genes are indicated by bold text.

